# Combining quantum cascade lasers and plasmonic metasurfaces to monitor *de novo* lipogenesis with vibrational contrast microscopy

**DOI:** 10.1101/2025.03.30.646207

**Authors:** Steven H. Huang, Dias Tulegenov, Gennady Shvets

## Abstract

The combination of a tunable quantum cascade laser (QCL) and plasmonic mid-infrared (MIR) metasurface is a powerful tool enabling label-free high-content microscopy of hydrated cells using the vibrational contrast of their constituent biomolecules. While the QCL provides a high-brightness source whose frequency can be rapidly tuned to that of the relevant molecular vibration, the metasurface is used to overcome water absorption of MIR light. Here we employ the resulting Metasurface-enabled Inverted Reflected-light Infrared Absorption Microscopy (MIRIAM) tool for non-destructive monitoring of the vital process of *de novo* lipogenesis (DNL), by which fat tissue cells (adipocytes) synthesize fatty acids from glucose and store them inside lipid droplets. Using ^13^C-labeled glucose as a metabolic probe, we produce spatially- and temporally-resolved images of ^13^C incorporation into lipids and proteins, observed as red-shifted vibrational peaks in the MIR spectra. These findings demonstrate MIRIAM’s capability for studying metabolic pathways with molecular specificity, offering a powerful platform for label-free imaging of cellular metabolism.

## Introduction

Among the many contributions of Federico Capasso to science, two especially stand out for the biophotonics community: the invention of the quantum cascade laser (QCL) [1] and the pioneering work in the area of mid-infrared (MIR) plasmonic metasurfaces [2].

The MIR spectral range is particularly important for life science applications because it overlaps with the vibrational frequencies of many chemical groups prevalent in biomolecules. For example, vibrational features within the 1,500 cm^−1^ < *ν* < 1,750 cm^−1^ MIR spectral window correspond to a combination of C=O stretching, N-H bending and C-N stretching in the amide backbone of proteins, as well as C=O stretching of the ester group in lipids [3]. Consequently, the ability to rapidly tune the emission wavelength (*λ* ≡ 1/*ν*) of a QCL to match various vibrational fingerprints of cellular molecular constituents (e.g., proteins, lipids, carbohydrates, and nucleic acids) has become highly attractive for label-free spectroscopy of cells and tissues, where the vibrational modes serve as endogenous image contrast [4]–[9].

Today, major manufacturers of MIR instrumentation (e.g., Agilent, Bruker, Daylight Solutions) have integrated QCLs into their MIR microscopes, offering QCL-driven discrete-frequency infrared (DFIR) microscopes on the market, and MIR micro-spectroscopy—using either broadband incoherent light sources for Fourier transform infrared (FTIR) spectroscopy or coherent ones, such as QCLs—has become a popular tool for analyzing biological samples [10]–[13]. Commercial DFIR microscopes have been used to analyze thin tissues section, dried biofluids, and fixed/dried cells [7], [8], [14], [15]. However, extending MIR spectroscopy to live (hydrated) cells and tissues has been hindered by the strong attenuation of MIR light in water. The use of thin flow cells and attenuated total reflection (ATR) geometries [16]–[20] have mitigated this issue to some extent but come with limitations, such as incompatibility with microwell-based cell analysis and automated liquid handling necessary for high-throughput sample analysis [21], [22].

To address these limitations, we have recently demonstrated that MIR plasmonic metasurfaces [23]–[28] supporting surface-enhanced infrared absorption (SEIRA) [29], [30] can be an effective tool to probe the vibrational spectra of living cell samples. Specifically, metasurfaces serve two essential functions: (i) they provide strongly localized near-field enhancements (within 100 nm from the surface) of optical fields, and (ii) they produce strong reflection signals that can be collected using an inverted infrared microscope operating in reflection mode (see Fig. 1). Cells are cultured on a metasurface comprising an array of plasmonic nanoantennas fabricated on an IR-transparent CaF2 substrate. The metasurface reflectance spectrum is modulated by interactions between the cells and the plasmonic near-field via SEIRA and these interactions are probed in the far field by measuring the reflectance spectrum. When combined with an FTIR spectrometer, these metasurfaces reveal rich molecular information and temporal dynamics about biomolecules in the plasma membrane and cytoskeleton.

**Figure 1.**
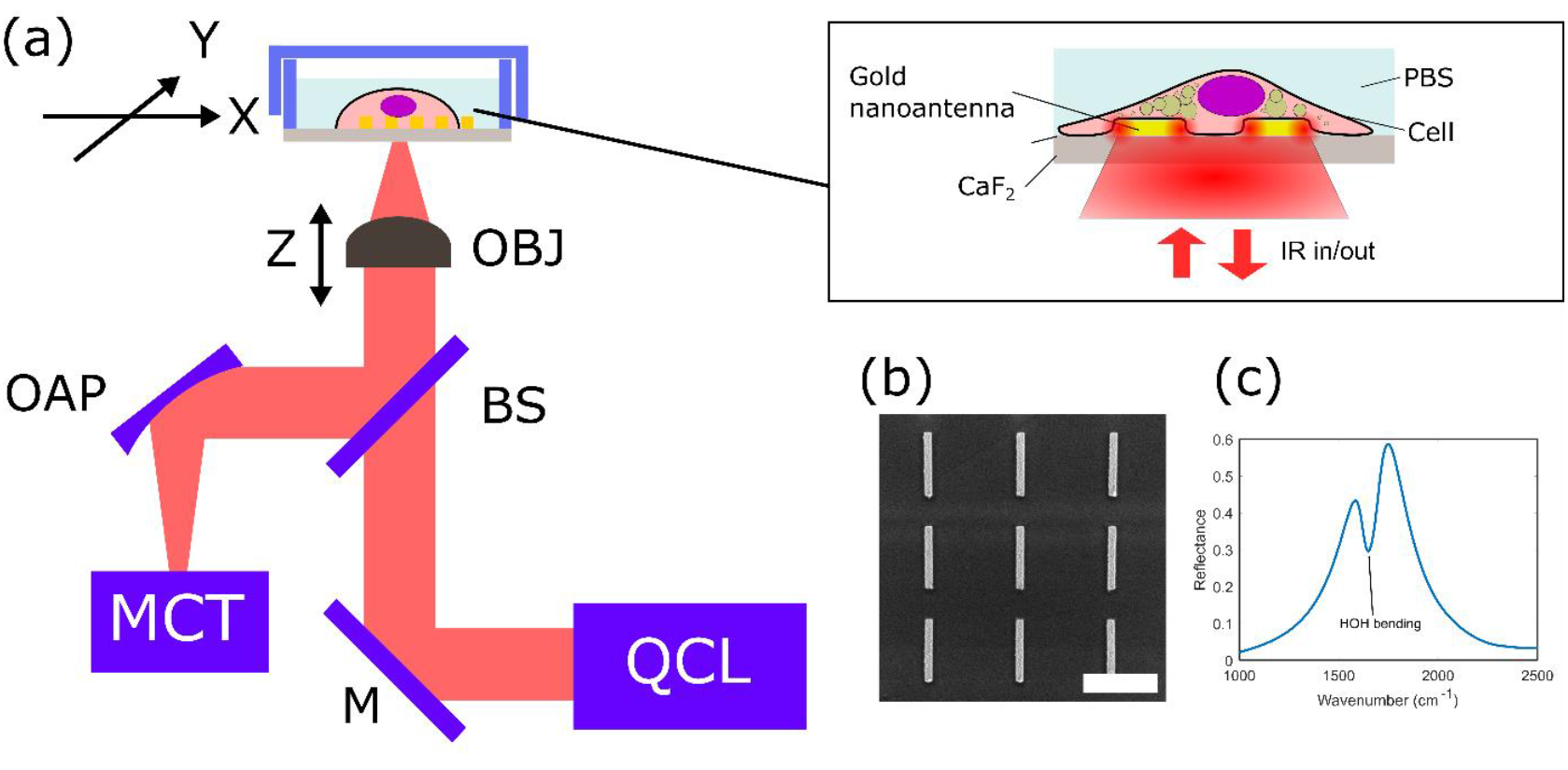
Metasurface-enabled Inverted Reflected-light Infrared Absorption Microscopy (MIRIAM). (a) Schematic drawing of the experimental setup. OBJ: objective, OAP: off-axis parabolic mirror, BS: beam splitter, M: mirror, MCT: mercury-cadmium-telluride IR detector, QCL: quantum cascade laser. (b) Scanning electron microscope image of the plasmonic nanoantennas. Scale bar: 2 µm. (c) Reflectance spectrum of the metasurface in water, measured by FTIR. The dip at 1650 cm^−1^ corresponds to H-O-H bending mode of water.

While our earlier work focused on FTIR-based MIR spectroscopy of live cells, effectively averaging the MIR spectra over many cells on top of the metasurface (typically a square area with several hundred micrometers in width), being able to image at single-cell resolution is highly desirable due to the inherent heterogeneity of live cells in physiologically relevant samples. To that end, we recently developed Metasurface-enabled Inverted Reflected-light Infrared Absorption Microscopy (MIRIAM), a hyperspectral platform for live-cell imaging with diffraction-limited resolution using QCL as the light source [31]. Previously, we demonstrated MIRIAM’s capabilities for vibrational microscopy of cells, detecting molecular contrasts from phosphates, proteins, and lipids, which allowed clear visualization of sub-cellular structures like nuclei and lipid droplets. Unlike conventional IR microscopy, MIRIAM overcomes the attenuation of IR light in water by confining the sensing volume to a few hundred nanometers around plasmonic hotspots. Additionally, the planar metasurface design and reflection-mode imaging facilitate straightforward integration with standard cell culture chambers, making MIRIAM suitable for high-throughput formats.

An especially promising application of MIRIAM is the development of a non-perturbative, all-optical, imaging platform for monitoring metabolic processes in cells (e.g., protein turnover, synthesis, or de novo lipogenesis) [32]–[38], as metabolomics provides a readout closest to the phenotype [39], [40]. Studying cellular metabolism presents unique challenges since small molecules like glucose and lipids cannot be labeled with fluorescent tags without disrupting their metabolic roles [39]. Stable isotope labeling with ^2^H and ^13^C, combined with analytical techniques like mass spectrometry or nuclear magnetic resonance, has become a widely used approach for metabolic studies [32], [41]–[44], However, these methods are inherently destructive and lack spatial resolution. Vibrational spectroscopy and imaging techniques, such as Raman microscopy, infrared (IR) microscopy, and mid-IR photoacoustic microscopy, have gained attention as non-destructive, spatially resolved tools for analyzing biomolecules labeled with stable isotopes [8], [38], [45]. These approaches enable label-free analysis with molecular specificity, and various labeling strategies—including ^13^C and ^2^H isotopes and probes with small azide or alkyne functional groups—have been developed for metabolic studies [8], [46]. Despite their promise, these techniques have limitations for cellular studies. For instance, Raman spectroscopy suffers from low signal intensity due to the small Raman cross-section of biomolecules, often necessitating high laser power, which can lead to photodamage and phototoxicity [47]. Similarly, mid-IR photothermal microscopy, while offering sub-500 nm spatial resolution comparable to optical microscopy, has a compromised signal-to-noise ratio when used with aqueous samples due to water’s small thermo-optic coefficient [45], [48], [49].

In this manuscript, we explore the application of the MIRIAM platform for characterizing metabolic processes, focusing on monitoring de novo lipogenesis (DNL) in adipocytes (fat tissue cells). DNL—the process by which cells synthesize fatty acids from non-lipid precursors like glucose—is a crucial pathway in adipocyte biology and systemic metabolic health [50], [51]. In adipocytes, DNL provides the basis for energy storage in the form of triglycerides, linking glucose metabolism to lipid synthesis via the tricarboxylic acid (TCA) cycle and subsequent lipogenic pathways. Dysregulation of DNL has been implicated in metabolic disorders such as obesity and type 2 diabetes, making adipocyte metabolism a promising target for cardiometabolic disease treatments.

We used MIRIAM to investigate DNL in mouse adipocyte cells using ^13^C-glucose as an IR-active probe. When substituted for ^12^C-glucose in the culture medium, ^13^C-glucose is metabolized into lipids stored in lipid droplets (LDs). The heavier ^13^C isotope induces a redshift in vibrational modes compared to ^12^C, detectable in IR spectra. We demonstrate that ^13^C incorporation into lipids and proteins is readily visualized with MIRIAM, reflecting glucose metabolism rates. Our findings highlight MIRIAM’s imaging capabilities and its sensitivity to ^13^C as a metabolic label, paving the way for parallel use of multiple metabolic labels to uncover intricate metabolic profiles through image-based cellular analyses.

## Experimental

### Metasurface Fabrication

The plasmonic nanoantennas were fabricated using electron beam lithography (EBL) on calcium fluoride (CaF_2_) substrate using PMMA resist. A 5 nm chromium adhesion layer and a 70 nm gold layer were deposited sequentially using thermal evaporation. The plasmonic nanoantennas were then formed by a lift-off process in acetone. The resulting metallic nanoantennas were arranged in an array covering a 500 µm × 500 µm area, constituting the metasurface. To enable imaging in aqueous environments, the metasurface was attached to the bottom of a cell culture chamber superstructure (CS16-CultureWell Removable Chambered Coverglass, Grace Bio-Labs).

### Cell Culture, Differentiation, and ^13^C Labeling

Undifferentiated mouse embryonic fibroblast 3T3-L1 cells (American Type Culture Collection) were cultured in pre-adipocyte expansion medium consisting of Dulbecco’s Modified Eagle Medium (DMEM, Gibco) with 4.5 g/L glucose and GlutaMAX supplement, 10% fetal bovine serum (FBS, Gibco), and 1% penicillin/streptomycin (P/S, Gibco). Cells were maintained at 37°C in a 5% CO2 incubator.

Upon reaching confluency, cells were incubated for another 48 hours before switching to differentiation medium. The differentiation medium consisted of DMEM supplemented with 10% FBS, 1% P/S, 1 μM dexamethasone (Sigma-Aldrich), 500 μM 3-isobutyl-1-methylxanthine (IBMX, Sigma-Aldrich), 2 μM rosiglitazone (Sigma-Aldrich), and 2 μg/mL bovine insulin (Sigma-Aldrich). [52] After 48 hours in differentiation medium, the cells were transitioned to adipocyte maintenance medium, composed of DMEM supplemented with 10% FBS, 1% P/S, and 2 μg/mL bovine insulin, and cultured for an additional 5 days.

Then, cells were harvested using 0.25% Trypsin-EDTA (Gibco) and re-seeded onto metasurfaces coated with 0.2% gelatin. Cells were divided into eight samples, each exposed to varying durations of ^13^C glucose maintenance medium and ^12^C glucose maintenance medium. Samples were labeled day 0 to day 7, corresponding to the number of days in ^13^C glucose maintenance medium. For the ^12^C glucose maintenance medium, regular DMEM was used, while for the ^13^C glucose maintenance medium, glucose-free DMEM was supplemented with 4.5 g/L U-^13^C_6_ glucose (Cambridge Isotope Laboratories) and the same concentrations of FBS, P/S, and insulin as in the ^12^C medium.

Seven days after seeding on the metasurfaces, cells were fixed with 4% formaldehyde for 15 minutes, washed three times with phosphate buffered saline (PBS), and stored in PBS prior to IR measurements.

### Inverted Mid-Infrared Microscope Setup

The inverted mid-infrared (MIR) microscope used in this study has been described in detail elsewhere. [31] Briefly, the MIR beam from a quantum cascade laser (QCL, MIRcat-1400, Daylight Solutions) was focused onto the metasurface from below using a Black Diamond-2 infrared collimation lens with a numerical aperture (NA) of 0.71 (390093, LightPath Technologies). The focal spot size was approximately 5 µm. The reflected beam from the metasurface was collected by the same objective lens and detected by a liquid-nitrogen-cooled mercury-cadmium-telluride (MCT) detector (MCT-13-1.0, InfraRed Associates).

The output current from the MCT detector was sent to a lock-in amplifier (SR830, Stanford Research Systems) for signal demodulation. For imaging, the metasurface sample was mounted on a dual-axis motorized microscope stage (HLD117, Prior Scientific), and point-scanning was performed by moving the sample through the laser focus. Hyperspectral image cubes were acquired by collecting discrete frequency images in the range of 1,500–1,800 cm^−1^ at 5 cm^−1^ intervals, with 2 µm pixel size.

To reduce water vapor absorption in the MIR, the entire optical setup was enclosed in an optical enclosure purged with dried air.

### Data Processing and Analysis

The acquired images were first processed using translational image registration in ImageJ to correct for sample drift and laser pointing fluctuations. Further data processing was performed in MATLAB. To correct for detector non-linearity, the signal intensity at each pixel was adjusted using a pre-measured calibration curve.

Fluctuations in laser intensity were accounted for by mean-centering each row of the image.

Absorbance was calculated using the formula:

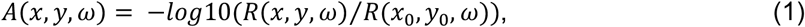

where *R*(*x, y, ω*) is the reflectance at a given pixel (*x, y*) and wavenumber *ω*, and *R*(*x*_0_, *y*_0_, *ω*) is the reflectance of a bare metasurface at a cell-free location. The hyperspectral image was subsequently de-noised using a minimum noise fraction (MNF) transform, retaining the first 20 components of the data for further analysis.

To quantify the intensity of specific vibrational peaks, a linear baseline subtraction method was applied to correct for shifts in the absorbance baseline caused by metasurface resonance variations. Two baseline points were selected on either side of each peak for the baseline correction. For the amide II peak, the absorbance at 1555 cm^−1^ was used to calculate the intensity, with baseline points at 1500 cm^−1^ and 1600 cm^−1^. For the ^13^C=O ester peak, the absorbance at 1705 cm^−1^ was used, with baseline points at 1685 cm^−1^ and 1720 cm^−1^. For the ^12^C=O ester peak, the absorbance at 1745 cm^−1^ was used, with baseline points at 1725 cm^−1^ and 1760 cm^−1^.

Additionally, the ^13^C=O ester peak exhibited significant overlap with the amide I peak of proteins and the H–O–H bending mode of water. [8], [45] To correct for these overlaps, the following model was used:

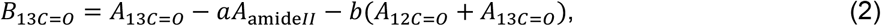

where *B*_13*C=0*_ is the apparent peak intensity calculated using the linear baseline method, and *A*_13*C=0*_, *A*_*amideII*_, and *A*_12*C=0*_ are the true peak intensities without spectral overlap. This model is based on two assumptions: (i) the spectral overlap in the amide II and ^12^C=O ester peaks is negligible; (ii) the overlap from the amide I peak is proportional to the number of amide groups whose combined bond strength is proportional to *A*_*amideII*_, and the overlap from the H–O–H bending mode arises from the total lipid-associated displacement of water proportional to (*A*_1*2C=0*_ + *A*_1*3C=0*_).

The coefficients *a* = 0.21 and *b* = 0.045 were empirically determined. The coefficient *a* was chosen to ensure that *A*_13*C*=*0*_ was zero in cytosolic regions lacking lipid droplets, while *b* was selected such that *A*_13*C*=*0*_ was zero in lipid droplets from the day 0 sample, which was cultured entirely in ^12^C glucose medium.

## Results

### MIRIAM-Based Imaging of Adipocyte Cells

Mid-infrared (MIR) hyperspectral images of 3T3-L1 adipocyte cells were acquired using MIRIAM, a platform previously developed by our group. [31] A schematic of the experimental setup is shown in Figure 1(a). The metasurface used in this study comprised an array of gold nanoantennas with dimensions of 200 nm in width, 1.8 µm in length, and 2.7 µm periodicity in both horizontal and vertical directions (Figure 1b).

When excited with incident light polarized along the long axis of the nanoantennas, the metasurface exhibited a resonance mode at approximately 1650 cm^−1^, with a pronounced peak in reflectance (Figure 1c). The resonance, characterized by a full-width at half maximum (FWHM) of 300 cm^−1^, covered multiple vibrational modes within the 1500–1800 cm^−1^ range. This single-resonance metasurface design was employed to probe key vibrational modes associated with adipocytes, including amide II (1555 cm^−1^) and amide I (1655 cm^−1^) for proteins, and ^13^C=O ester (1705 cm^−1^) and ^12^C=O ester (1745 cm^−1^) for lipids.

A representative bright-field image of fixed 3T3-L1 adipocyte cells on the metasurface is shown in Figure 2(a). Lipid droplets (LDs) within the cells are clearly visible due to the high refractive index contrast between lipids and water. The gold nanoantennas of the metasurface appear as an opaque periodic pattern in the background. Figures 2(b)-(d) display the raw signal, reflectance spectra, and absorbance spectra, respectively, for the LD and cytoplasm regions.

**Figure 2.**
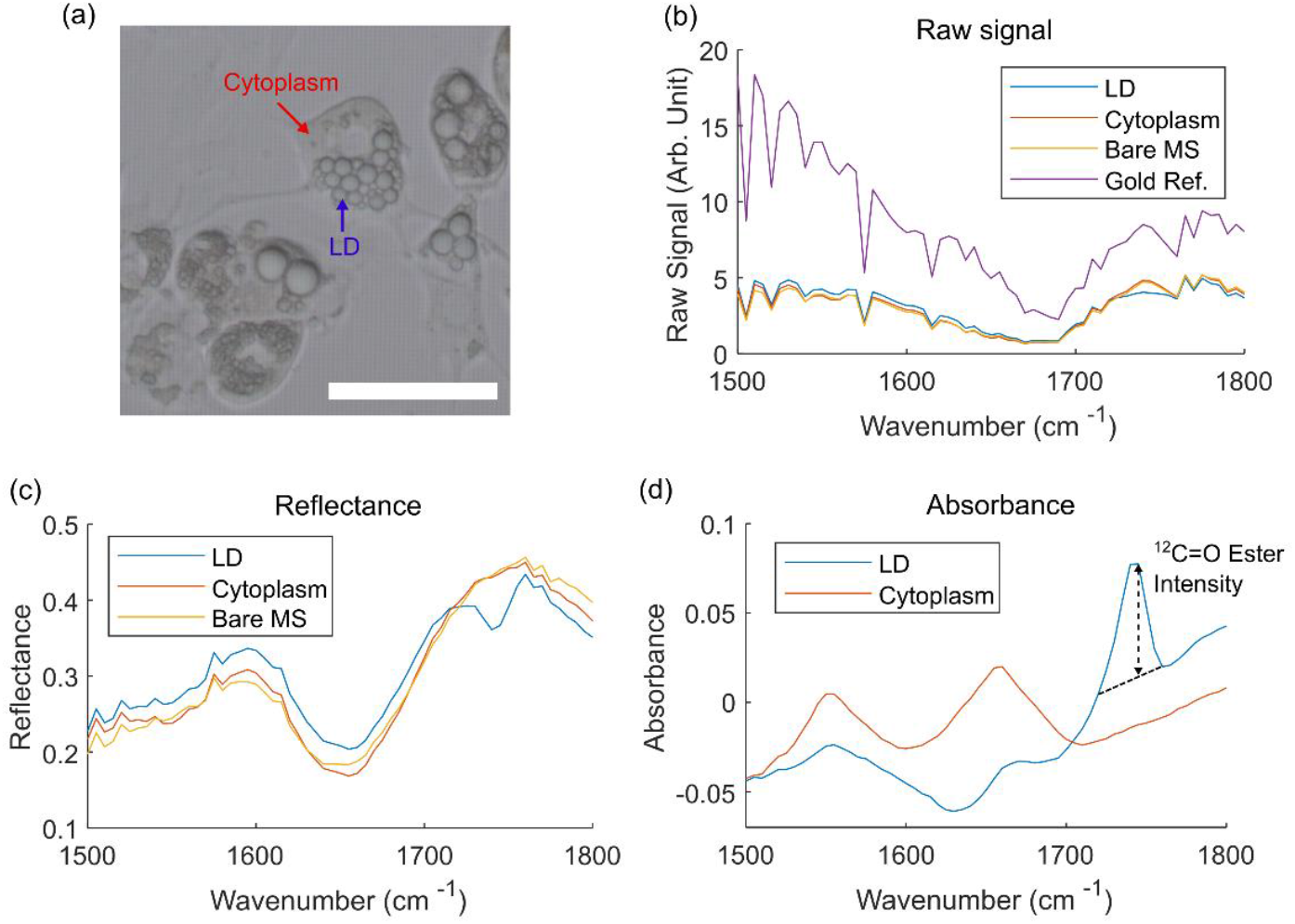
MIR spectra of 3T3-L1 adipocyte acquired through MIRIAM. (a) Bright-field image of 3T3-L1 adipocytes on the metasurface, with the periodic nanoantenna pattern visible in the background. Scale bar: 100 µm. (b) Raw reflection signal at different locations. “Bare MS” refers to a metasurface region without cells, while “Gold Ref.” indicates a gold patch used for reflectance reference measurements. Sharp peaks in the spectra correspond to atmospheric water vapor absorption. (c) Reflectance spectra at different positions. (d) Absorbance spectra at LD and cytoplasm position. The black dotted line shows the linear baseline correction used to determine the ^12^C=O ester intensity.

The intensity variation in the raw signal spectra (Figure 2(b)) results from laser intensity fluctuations with respect to wavenumber and atmospheric water vapor absorption. This variation is normalized by collecting a reference spectrum from a 70 nm thick gold patch adjacent to the metasurface, yielding the reflectance spectra shown in Figure 2(c).

Although our QCL-based measurement lacks the spectral range to fully capture the metasurface’s resonance mode (see FTIR measurement in Figure 1(c) for the full reflectance spectrum), variations in reflectance due to the metasurface resonance and a dip corresponding to the H-O-H bending mode of water are evident.

To isolate the vibrational signature of the cells, absorbance spectra are calculated using a reference spectrum collected from a bare metasurface position (Equation 1). The absorbance spectra reveal vibrational modes corresponding to cellular biomolecules, including the amide I (1655 cm^−1^) and amide II (1555 cm^−1^) modes from cytoplasmic proteins, as well as the ^12^C=O ester mode (1745 cm^−1^) from lipids in the LDs (Figure 2d). In MIRIAM, the vibrational signals of analytes are superimposed on the metasurface resonance spectrum. This results in a varying baseline in the absorbance spectrum due to shifts in the metasurface resonance when analytes interact with the near-field. To accurately quantify vibrational signals, a linear baseline subtraction was applied, as described in the Experimental section. Figure 2(d) shows the baseline subtraction for ^12^C=O ester mode, as an example.

Although previous studies have demonstrated MIRIAM’s capability to image living cells at a few discrete frequencies, [31] the current acquisition speed of ~5 minutes for a 500 µm × 500 µm image makes hyperspectral imaging of living cells impractical, especially when collecting data for 10 or more frequencies. Movement of living cells during the measurement introduces spectral inaccuracies, particularly for small structures like lipid droplets. To address this limitation, this study focused on imaging fixed cells while keeping the cells hydrated in PBS to avoid disruptions in cellular structures and morphology.

To study the glucose metabolism and de novo lipogenesis in 3T3-L1 adipocytes, eight samples of 3T3-L1 adipocyte cells were prepared with varying exposure times to ^13^C glucose (Figure 3). Preadipocyte cells were chemically induced to differentiate into adipocytes and cultured for 5 days in a flask until lipid droplets were visibly formed. At this point, the cells were reseeded onto metasurfaces and incubated in maintenance media with differing exposure durations to ^13^C glucose (0–7 days). This design ensured similar lipid droplet maturation times across samples, with comparable droplet sizes, while varying the relative amounts of ^13^C=O ester and ^12^C=O ester in lipid droplets due to differential exposure to ^13^C glucose and ^12^C glucose.

**Figure 3.**
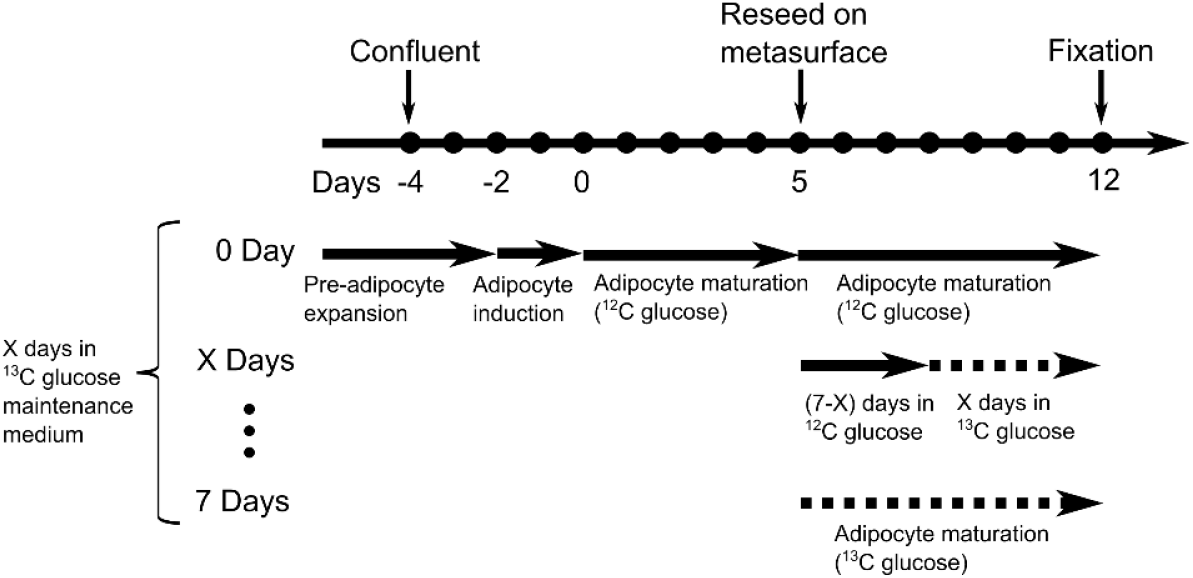
Experimental timeline for differentiation and ^13^C glucose feeding of 3T3-L1 adipocytes.

Representative MIRIAM-based MIR images of samples with 0 days (no exposure) and 7 days of ^13^C glucose exposure are shown in Figure 4. The amide II channel, derived from the protein-associated vibrational contrast, exhibits distinctly different spatial distribution compared to the ^13^C=O and ^12^C=O ester channels derived from the lipids-associated contrast. This demonstrates the specificity of MIRIAM for detecting distinct molecular vibrations with minimal crosstalk between channels. Owing to the limited penetration depth (~100 nm) of the plasmonic near-field surrounding the antennas, the protein signal predominantly originates from the focal adhesion sites and the cytoskeletal layer around the cell cortex. Protein contrast highlights cell spreading on the metasurface, with individual cells clearly discernible.

**Figure 4.**
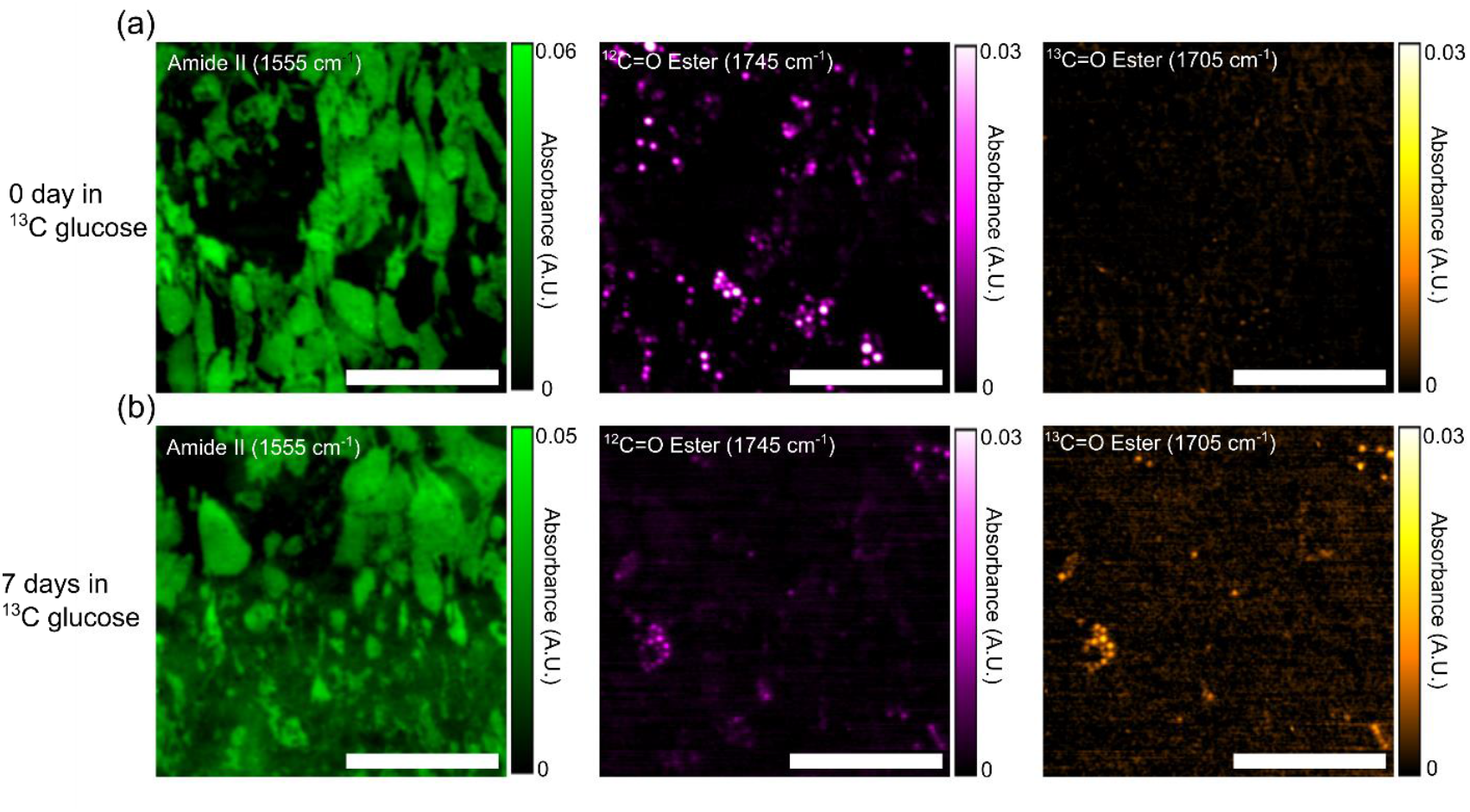
Fixed 3T3-L1 adipocyte IR absorbance images acquired by MIRIAM. (a) 0 day exposure to ^13^C glucose and (b) 7 days exposure to ^13^C glucose. Increase in ^13^C=O ester and decrease ^12^C=O ester in the 7 days sample can be clearly seen. Scale bar: 200 µm.

In contrast, the lipid channel corresponds to lipid droplets. However, many lipid droplets visible in bright-field microscopy (Figure 2a) are not detected in the MIR images. This is attributed to the surface sensitivity of MIRIAM, which renders lipid droplets positioned far from the metasurface essentially invisible. Despite this limitation, sufficient lipid droplets were visible to quantify ^12^C=O and ^13^C=O ester signals.

In the 0-day sample, lipid droplets exhibited no detectable ^13^C=O ester signal, as expected, with all lipid derived from ^12^C glucose. In contrast, the 7-day sample showed a marked increase in the ^13^C=O ester signal and a corresponding decrease in the ^12^C=O ester signal within lipid droplets. The colocalization of ^13^C=O and ^12^C=O signals confirmed that both signals originated from the same lipid droplets, with the ratio of the two signals reflecting the duration of incorporation of ^13^C into the lipid droplets.

We note that the combination of the inverted QCL-based microscope and the metasurface enabled diffraction-limited imaging of sub-cellular LDs inside hydrated cells. Earlier attempts at spectrochemical imaging using commercial widefield MIR microscopes worked exclusively with dried samples and encountered coherence-related speckle artifacts in transmission-mode images [7], [8]. By using a plasmonic metasurface in addition to a widefield MIR microscope, Rosas *et al*. demonstrated enhanced MIR spectra of dried tissue samples in both transmission and reflection mode, but no applications to hydrated samples were reported [9]. Likewise, confocal MIR microscopy without a metasurface produced speckle-free vibrational images with diffraction-limited resolution, but only in the transmission mode [5], [6], which cannot be used for imaging hydrated cells.

### Tracking ^13^C incorporation in lipids and proteins through IR spectra

To investigate the temporal evolution of ^13^C glucose metabolism, representative spectra were extracted from each sample (Figure 5), focusing on two distinct regions: lipid droplets (Figure 5a) and the cytoplasmic region outside of the lipid droplets (Figure 5b).

**Figure 5.**
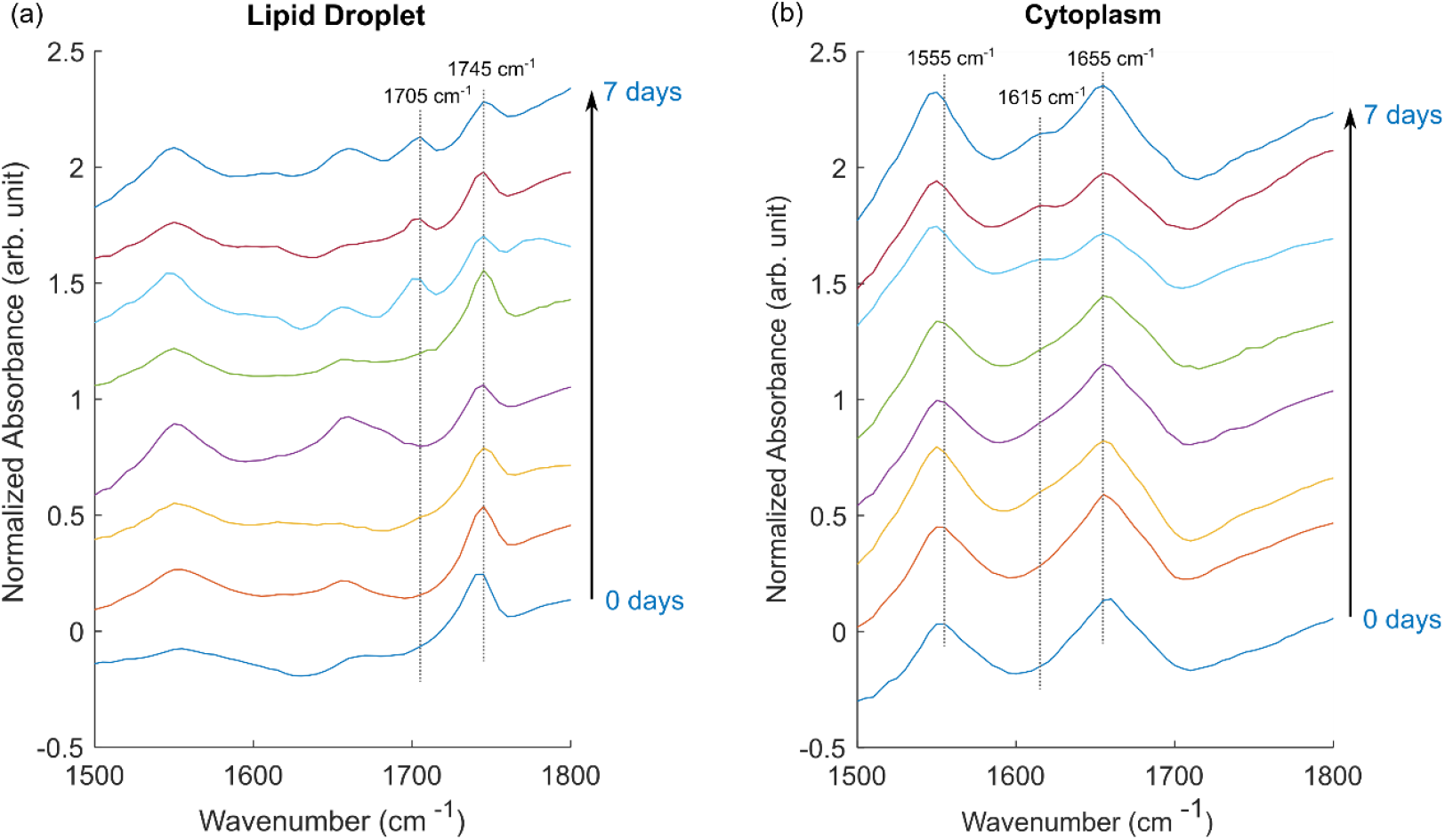
Representative IR spectra from (a) lipid droplets and (b) cytoplasm regions without lipid droplets, for samples with varying exposure times to ^13^C glucose. An increase in the ^13^C=O ester peak (1705 cm^−1^) and the ^13^C amide I peak (1615 cm^−1^) is observed with longer exposure to ^13^C glucose medium.

In the lipid droplet spectra, increasing exposure time to ^13^C glucose led to a pronounced increase in the ^13^C=O ester peak at 1705 cm^−1^ and a corresponding decrease in the ^12^C=O ester peak at 1745 cm^−1^. This shift was most prominent in the samples with 5–7 days of exposure. Additionally, all spectra displayed variable but significant amide I and amide II peaks, likely attributed to the protein from cell cortex between the lipid droplet and the metasurface. The irregular shape of the amide I peak at 1655 cm-1 is partially explained by its overlap with the H-O-H bending mode of water. Lipid droplets displace water in their vicinity, resulting in reduced absorbance from the H-O-H bending mode relative to the bare metasurface used for background measurements.

In the cytoplasmic spectra (Figure 5b), the amide I and II peaks dominate, as expected. However, in samples with the highest ^13^C incorporation (5–7 days), a small peak emerges at 1615 cm^−1^, attributed to the ^13^C amide I peak. [8], [53] Additionally, the amide II peak exhibits a slight shift of approximately 5 cm^−1^ in these samples, also reflecting the incorporation of ^13^C into amino acids. These observations suggest substantial *de novo* amino acid biosynthesis using glucose as the carbon source.

Glucose metabolism is known to be elevated in adipocyte cells, consistent with their role in lipid storage. [54] Our findings indicate that carbon from glucose metabolism is utilized not only for *de novo* lipogenesis but also for the synthesis of amino acids. This likely involves the tricarboxylic acid (TCA) cycle, which provides precursors such as α-ketoglutarate for glutamate and oxaloacetate for aspartate. [55] This coupling between the *de novo* synthesis of fatty acids and amino acids highlights a metabolic interdependence that warrants further investigation.

### Temporal dynamics of de novo lipogenesis

To evaluate the temporal dynamics of de novo lipogenesis, we quantified the rate of lipid synthesis by calculating the ratio of the ^13^C=O peak intensity to the total lipid content (defined as the sum of the ^12^C=O and ^13^C=O peak intensities). A threshold mask was applied to the ^12^C=O image to identify lipid droplets (LDs) with a strong ^12^C signal, and the ratio was calculated using only the selected pixels. Approximately 100 pixels were analyzed per sample, representing 5–10 lipid droplets per sample. The results are presented in Figure 6.

**Figure 6.**
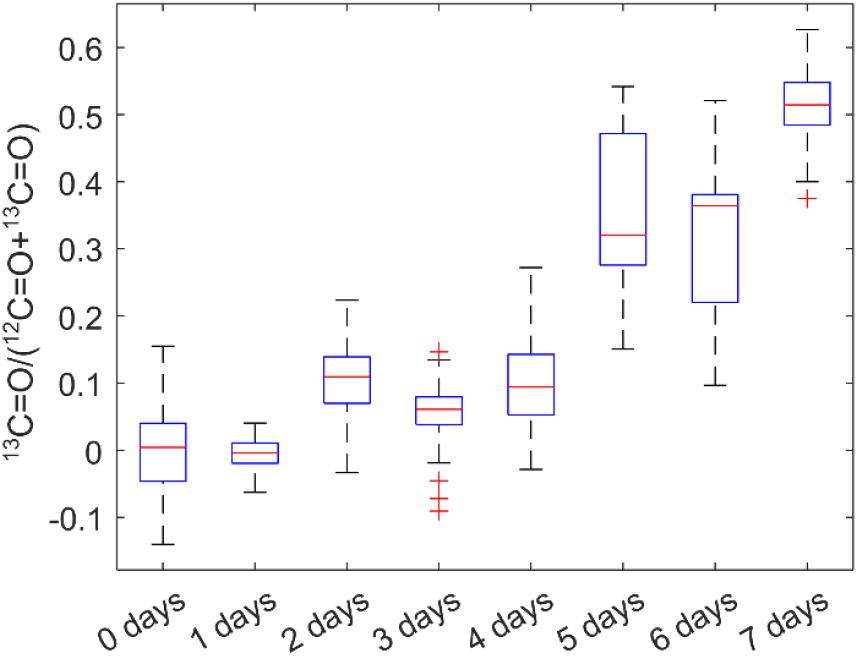
Ratio of ^13^C-labeled lipids to total lipids (^12^C + ^13^C) as a function of time. The intensities of ^13^C and ^12^C lipids were extracted from MIRIAM images, using approximately 100 pixels per sample, selected from regions with the highest ^12^C=O ester intensity. These pixels represent 5–10 lipid droplets per sample.

The ratio of ^13^C=O to total lipids gradually increased over time, reaching a value of 0.51 ± 0.05 after 7 days of exposure to ^13^C glucose. This ratio is slightly higher than reported in a previous study by Shuster et al. on de novo lipogenesis in 3T3-L1 cells, [45] which may be attributed to the longer exposure times used in our study (up to 7 days) compared to the 3-day exposure in the earlier study. Even though the cells have been exposed to ^13^C glucose for a long time (7 days), there is still significant amount of ^12^C=O in the lipid droplet. This can be explained by alternate carbon sources that the cells may be using for de novo lipogenesis, which include glutamine and free fatty acids, [43], [54] which are not labelled with ^13^C.

Interestingly, the rate of increase in the ^13^C=O ratio appeared to slow down after the first three days. For example, samples with 1–4 days of ^13^C glucose exposure exhibited significantly lower ^13^C=O ratios compared to samples with 5–7 days of exposure. This observation may be explained by a decline in the rate of de novo lipogenesis as lipid droplets mature over time. Alternatively, the observed slowing could be due to an accumulation of ^12^C=O lipids. As the lipid droplet size increases, the rate of change in the ^13^C=O-to-total-lipid ratio is expected to decrease even if the absolute rate of ^13^C=O generation remains constant. However, our technique’s limited penetration depth, determined by the plasmonic hotspot, prevents accurate measurement of individual lipid droplet sizes, and this hypothesis warrants further investigation.

We also observed considerable variability in the ^13^C=O-to-total-lipid ratio across the samples. While Shuster et al. attributed this variability to spatial heterogeneity in 13C incorporation, [45] we suggest that in our case, this spread is more likely due to experimental error in quantifying the ^13^C ratio. Similar variability was observed across samples with different ^13^C glucose exposure times, which suggests that the variation is not driven by biological differences. Potential sources of error include laser intensity fluctuations, environmental water vapor variability, or inaccuracies in quantifying the ^12^C=O and ^13^C=O intensities caused by spectral artifacts and distortions. Further refinements in experimental design and signal processing may help address these limitations.

## Conclusion

In this work, we utilized MIRIAM, a mid-infrared microscopy technique based on plasmonic metasurfaces, to image and monitor DNL in ^13^C-labeled 3T3-L1 adipocyte cells. ^13^C=O, introduced through ^13^C-labeled glucose, served as a metabolic label for DNL. We demonstrated the incorporation of ^13^C into both proteins and lipids by observing shifts in the amide I/II and C=O vibrational peaks due to ^13^C substitution. As expected, prolonged exposure to ^13^C glucose resulted in greater incorporation of ^13^C=O into lipid droplets.

While glucose metabolism is elevated in adipocyte cells, it is not the sole carbon source for anabolic processes. Other metabolites, such as glutamine and free fatty acids, are also critical contributors to biosynthetic pathways. In this study, only glucose was labeled with ^13^C, providing insights into its role in lipogenesis and protein synthesis.

Future studies could expand on this approach by labeling multiple metabolic precursors with IR-active probes, such as *2*H or azide-containing compounds. This would enable a more comprehensive elucidation of metabolic pathways and provide insights into the interplay between glucose metabolism and alternative anabolic substrates. Such advancements could significantly enhance our understanding of cellular metabolism and its regulation.

## Research funding

The research reported here was supported by the National Cancer Institute of the National Institutes of Health under award number R21 CA251052 and by the National Institute of General Medical Sciences of the National Institutes of Health under award number R21 GM138947. This work was performed in part at the Cornell NanoScale Facility, a member of the National Nanotechnology Coordinated Infrastructure (NNCI), which is supported by the National Science Foundation (Grant NNCI-2025233).

## Author contribution

All authors have accepted responsibility for the entire content of this manuscript and consented to its submission to the journal, reviewed all the results and approved the final version of the manuscript. SHH and GS designed the experiment. SHH and DT performed the experiment. SHH analysed the data. GS supervised this work. SHH prepared the manuscript with contributions from all co-authors.

## Data availability statement

Data for this study are available from the corresponding authors on reasonable request.

